# GHB degradation via TCA cycle is the major metabolic route in *Arabidopsis thaliana*

**DOI:** 10.1101/2022.11.04.515263

**Authors:** Dereje W. Mekonnen

## Abstract

Gamma-hydroxybutyric acid (GHB) is a reduced product of the chemically reactive succinic semi-aldehyde (SSA), which itself is produced from the degradation of GABA. It is regarded as a short chain fatty acid. Despite our understanding of the synthesis of GHB, little is known about its fate. Exogenous application of 0.1 mM GHB to *pop2 x ssadh*, a double mutant incapable of catabolizing GABA, increased the endogenous GHB level by 13-fold compared to the wild type. The GHB dynamic studies by feeding and relieving from treatments showed that GHB is rapidly metabolized in wild type plants compared to the *pop2 x ssadh* double mutant. Although regarded as a short chain fatty acid, GHB level was not altered in mutants of the beta oxidation pathway following exogenous feeding. Therefore, the metabolism back to SSA and then TCA cycle appears to be the major route for GHB degradation. However, the presence of another catabolic route such as secondary modifications cannot be ruled out.

---

The presence of GHB has been reported in various organisms including mammals (Giaman and Roth, 1964), yeast (Breitkreuz et al. 2003; Bach et al. 2009) and plants (Allan et al. 2003; Allan et al. 2008). The accumulation of GHB in *ssadh* mutants has been considered as the cause of the *ssadh* aberrant phenotype (Fait et al. 2004). In contrast, the accumulation of GHB in Arabidopsis plants exposed to environmental stress was regarded as a positive response (Allan et al. 2008). Our previous work resolved those contrasting views by feeding exogenous GHB to Arabidopsis and yeast (Mekonnen and Ludewig, 2016). The specific effect of GHB in Arabidopsis could not be resolved since its conversion back to SSA is active. In yeast, where there is no GHB to SSA activity, high concentrations of GHB did not elicit toxicity (Mekonnen and Ludewig, 2016). Therefore, the toxicity induced by high exogenous concentration of GHB in *Arabidopsis* plants is likely due to its conversion to a chemically reactive SSA.

GHB is produced from the reduction of SSA by the activity of GHB dehydrogenase (GLYR). In Arabidopsis, two GLYR genes have been reported previously (Simpson et al. 2008; Hoover et al. 2007). Enzymes encoded by these two genes catalyze only the forward reaction i.e. the conversion of SSA to GHB (Simpson et al. 2008; Hoover et al. 2007). Indeed, external feeding of GHB to the double knockouts (*glyr1/2*) did not show an altered phenotype compared to the wild type (Mekonnen and Ludewig, 2016). Although GHB synthesis is fairly studied in Arabidopsis, little consideration is given to its subsequent metabolism. To examine whether GHB metabolism back to the TCA cycle is the major route, and whether peroxisomal beta-oxidation is involved in GHB catabolism, mutant lines were characterized. Arabidopsis *pop2 x ssadh* double mutant accumulated 3-fold higher amounts of GHB under control conditions and 13-fold higher amounts when treated with 0.1 mM GHB than the wild type (Fig. 1A).. This observation suggests that GABA-T and SSADH are involved in the metabolism of GHB. Indeed, the involvement of GABA-T and SSADH in GHB metabolism has recently been suggested after succinate and GABA were induced following GHB treatment (Mekonnen and Ludewig, 2016). Interestingly, in *pop2 x ssadh* double mutants the GABA content was unchanged after GHB treatment (Fig. 1B), further confirming that GABA-T plays a role in GHB metabolism. However, the presence of more GHB in the double mutant under control condition is unexpected. Arabidopsis genome contains a single gene that encodes GABA-T (van Cauwanberghe et al. 2002). Knockout of this single gene is expected to shut off the pathway and hence prevent the accumulation of GHB. Nevertheless, considerable amount of GHB was detected in the *pop2 x ssadh* double mutant perhaps suggesting the existence of another enzyme with a transaminase activity. Clark et al. (2009) detected a small 2-oxoglutarate dependent GABA-T activity in crude protein extracts which was not detected in either truncated or full length GABA-T recombinant proteins. These observations might suggest the existence of a second 2-oxoglutarate-dependent low affinity GABA-T in Arabidopsis genome.

**Fig. 1.**
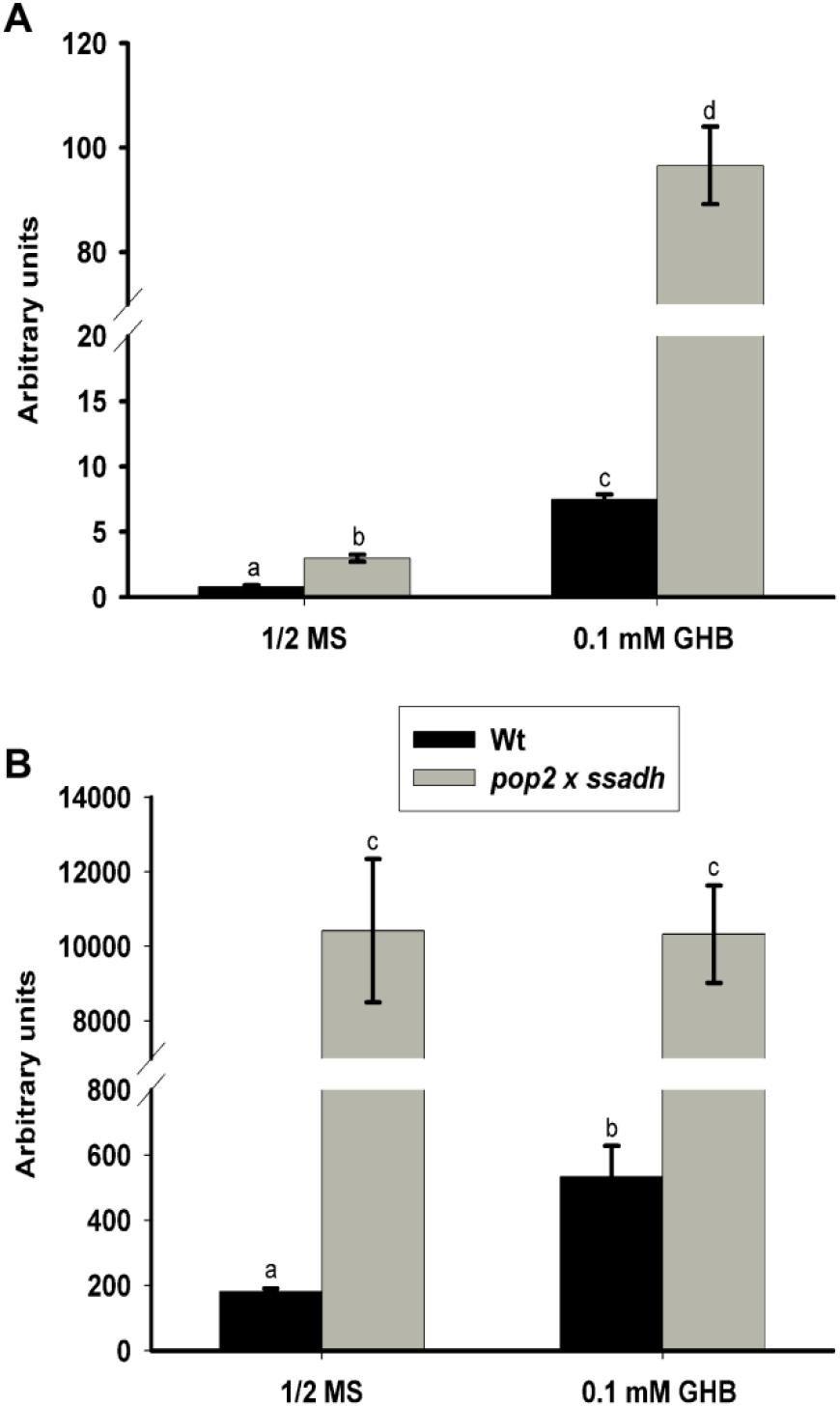
Analysis of GHB (A) and GABA (B) content in Wt and *pop2 x ssadh* mutant; seeds of Wt and *pop2 x ssadh* mutant were germinated and grown on ½ MS medium without and with 0.1 mM GHB; After two weeks of growth, samples were collected and snap-frozen in liquid nitrogen; 3 to 4 plants were pooled together to make one sample for *pop2 x ssadh* mutant; metabolite extraction and measurement was carried out as described previously;^7^ values are means of five biological samples; error bars represent the standard error of means; different letters show statistical significance after student’s *t*-test.

Furthermore, when 0.1 mM GHB treated *pop2 x ssadh* plants were transferred to GHB free medium, only 48% of the GHB was metabolized within 3 hours (Fig. 2B). In contrast, 72% of the GHB was metabolized in wild type after 3 hours of incubation on GHB free ½ MS plate (Fig. 2A). After 12 hours of incubation on GHB free medium, the amount of GHB detected in the tissue of the Wt was similar to the GHB amount in untreated samples; whereas, in the double mutant it was significantly higher (Fig. 1B). These observations suggest that; first GHB has short half-life time. i.e. it is rapidly metabolized than being stored; second, GABA-T and SSADH are involved in GHB metabolism. Poldrugo and Addolorato (1999) mentioned a short half-life of GHB once it is formed in the brain. The reduction of the GHB level in *pop2 x ssadh* tissue with increasing time on GHB free medium might suggest the presence of another route for the metabolism of GHB. GHB is regarded as short chain fatty acids and its metabolism via *β*-oxidative pathway to 3,4-dihydroxybutyrate (DHB) has been suggested in mammals (Poldrugo and Addolorato 1999; Jacobs et al. 1981). In mammals, *β*-oxidation occurs both in mitochondria and peroxisome; however, in plants it is regarded exclusively as peroxisomal (Poirier et al. 2006). However, Masterson and Wood (2001) showed the existence of a mitochondrial fatty acid *β*-oxidation which was active at certain stage of seedling development. To examine the degradation of GHB via *β*-oxidation, Arabidopsis *pxa1* and *kat2* mutants defective in peroxisomal *β*-oxidation were treated with GHB. *PXA1* encodes a peroxisomal ATP-binding cassette transporter, and is known to transport fatty acids into peroxisomes (Zolman et al., 2001). *KAT2* encodes a 3-ketoacyl-CoA thiolase that catalyzes the last step of the beta oxidation of fatty acids in peroxisomes (Footitt et al. 2007; Jiang et al. 2011). Both mutants, *pxa1* and *kat2*, were grown on 1.5 mM GHB containing plates for two weeks. Then, a portion of plants were transferred to GHB free plates. After two days of incubation on GHB free plates, almost nothing was left in the plant (Fig. 3), suggesting that GHB is not metabolized *via* peroxisomal beta oxidation pathway. The conversion back to SSA seems to be the major route for GHB metabolism. However, the metabolism of GHB in *pop2 x ssadh* mutant suggests the existence of another route for GHB degradation.

**Fig. 2.**
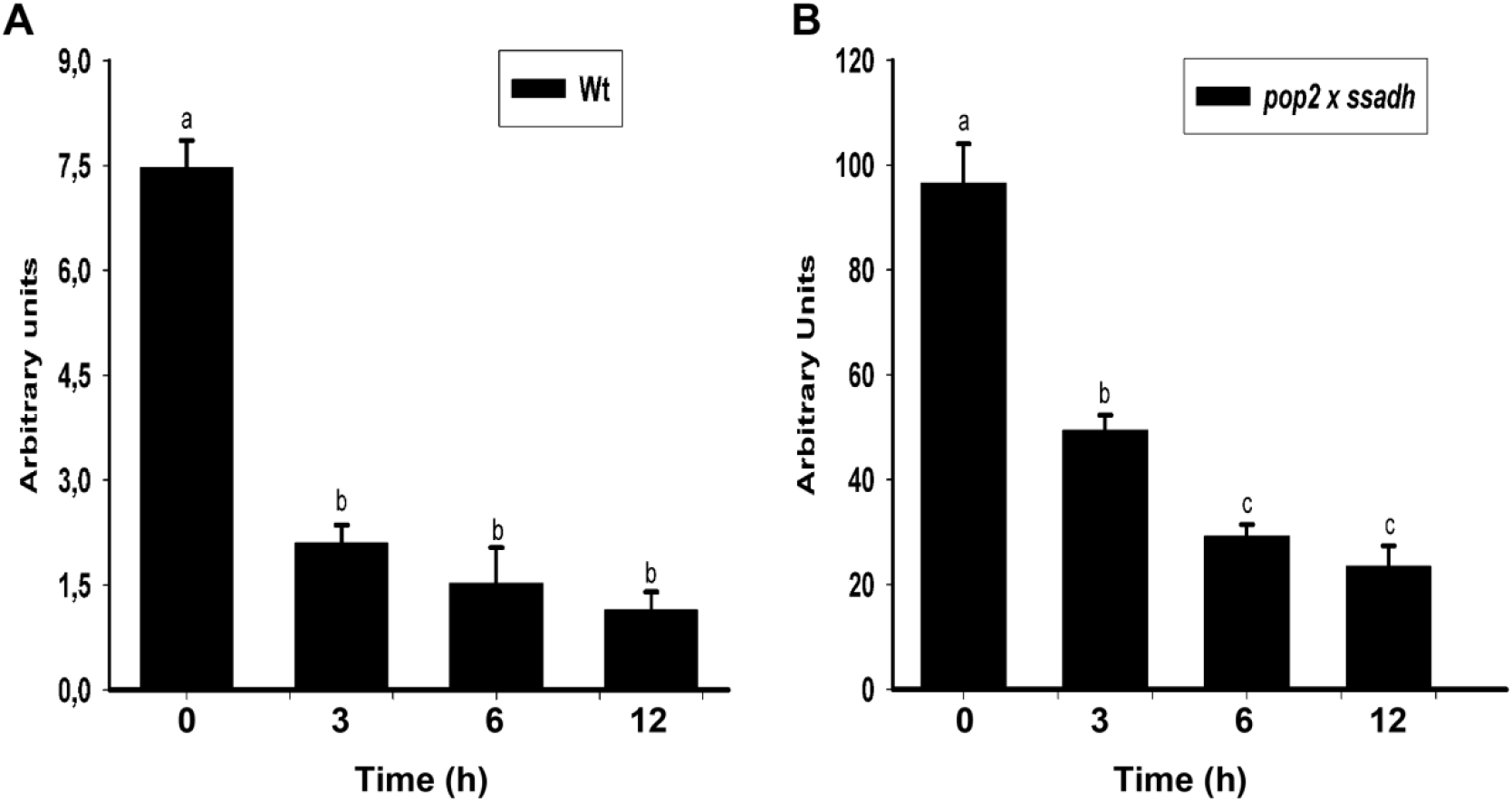
Analysis of GHB dynamics in the Wt (A) and *pop2 x ssadh* (B) lines; seeds of Wt and *pop2 x ssadh* were germinated and grown on ½ MS plate without and with 0.1 mM GHB; after two weeks, plants were transferred to GHB-free plates; samples were collected after 3, 6 and 12 hours for GC/MS analysis; metabolite extraction and measurement was carried out as described previously;^7^ values are means of five biological replicates; error bars represent the standard error of means; different letters show statistical significance after student’s *t*-test.

**Fig. 3.**
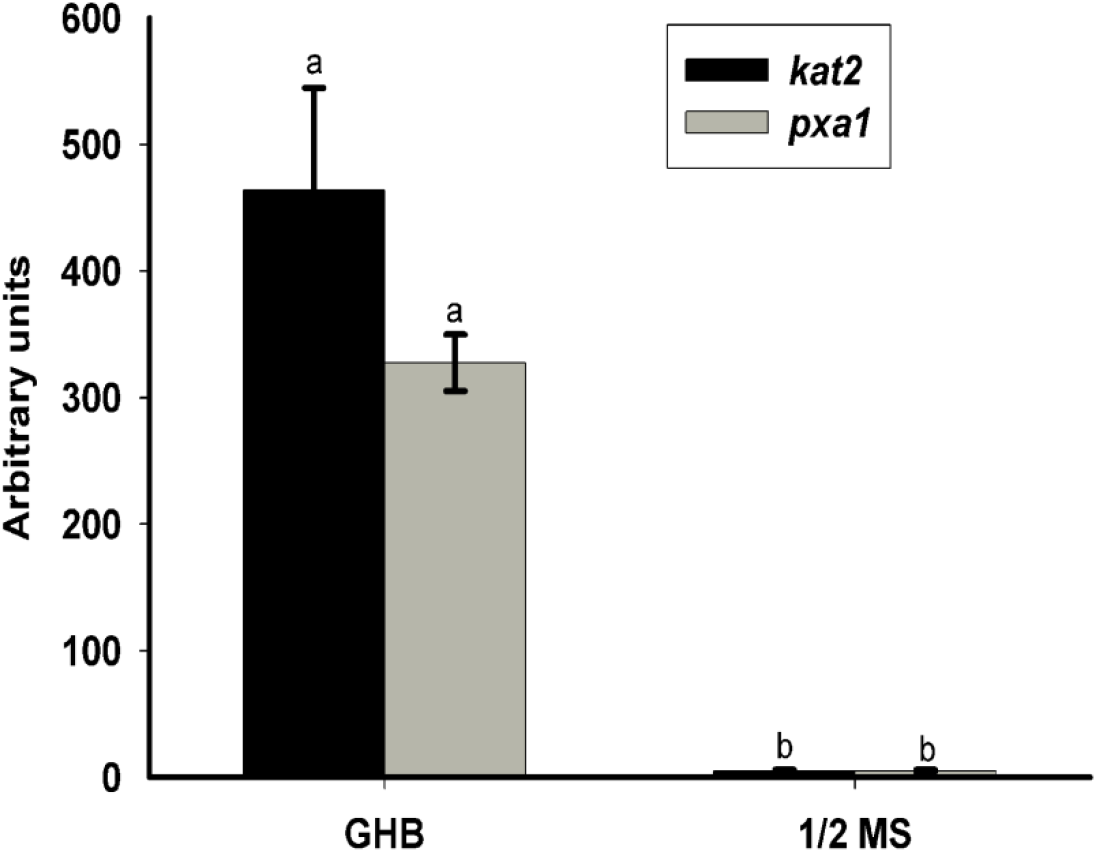
Analysis of GHB dynamics in *kat2* and *pxa1* mutants; seeds of *kat2* and *pxa1* were germinated and grown on ½ MS plate containing 1.5 mM GHB; after two weeks of growth some of the plants were transferred to GHB-free ½ MS plates; after two days on GHB free plates samples were collected for GHB measurement; extraction and measurement of GHB was carried out as described previously;^7^ values are means of five biological replicates; error bars show the standard error of means; different letters show statistical significance after student’s *t*-test.

## Future perspectives

In humans, the accumulation of GHB leads to a pathological disorder called GHB aciduria. As a medically important compound, understanding of the metabolic routes of GHB helps to identify more drugs targeting the enzymatic components involved in the metabolism. For that, metabolic analysis on a broader scale is needed to identify possible GHB derived metabolites. Amino acid derived metabolites undergo various modifications for storage or deactivation purposes. One of such modifications is glycosylation. Glycosylation involves the addition of activated sugar to the hydroxyl and carboxyl group of a compound. For example, isoleucic acid (ILA), a catabolic product of isoleucine metabolism, and N-hydroxypepicolic acid (NHP), a catabolic product of lysine, are glycosylated when they are in excess in the tissue (Bauer et al.,2021A; Bauer et al., 2021B). Glycosylation of GHB to its carboxylic and hydroxyl group can be considered as a possible route for a rapid metabolism of GHB when they are in excess in the tissue. Of course, the enzymes catalyzing GHB glycosylation will also be a subject of further investigation.

## Acknowledgement

I would like to thank the NRW IGSDHD program for funding this work.

## References

1. Giarman NJ, Roth RH. Differential estimation of gammabutyrolactone and gamma hydroxylbutyric acid in rat blood and brain. Science. 1964; 145:583–584.

2. Bauer S, Mekonnen DW, Geist B, Lange B, Zhang W, Schäffner AR. The Isoleucid acid triad distinct impats on plant defense, root growth and formation of reactive oxygen species. J. Exp. Bot. 2020a; 71(14):4258–4270.

3. Bauer S, Mekonnen DW, Hartmann M, Janowski R, Zhang W, Lange B, Geist B, Zeier J, Schäffner AR. UGT76B1, a promiscuous hub of small molecule-based immune signaling, glucosylates N-hydroxypipecolic acid and controls basal pathogen defense. Plant Cell. 2021b; 33:714–734.

4. Breitkreuz KE, Allan WL, Van Cauwenberghe OR, Jakobs C, Talibi D, Andre B, Shelp BJ. A novel c–hydroxylbutyrate dehydrogenase: identification and expression of an Arabidopsis cDNA and potential role underoxygen deficiency. J Biol Chem. 2003; 278:41552–41556.

5. Bach B, Meudec E, Lepoutre JP, Rossignol T, Blondin B, Dequin S, Camarasa C. New insights into gamma-aminobutyric acid catabolism: evidence for gammahydroxybutyric acid and polyhydroxybutyrate synthesis in Saccharomyces cerevisiae. Appl. Environ. Microbiol. 2009; 75:4231–4239.

6. Allan WL, Peiris C, Bown AW, Shelp BJ. Gamma-hydroxybutyrate accumulates in green tea leaves and soybean sprouts in response to oxygen deficiency. Can J Plant Sci. 2003; 83:951–953.

7. Allan WL, Simpson JP, Clark SM, Shelp BJ. γ-Hydroxybutyrate accumulation in Arabidopsis and tobacco plants is a general response to abiotic stress: putative regulation by redox balance and glyoxylate reductase isoforms. J. Exp. Bot. 2008; 59(9):2555–2564.

8. Fait A, Yellin A, Fromm H. GABA shunt deficiencies and accumulation of reactive oxygen intermediates: insight from Arabidopsis mutants. FEBS Letter. 2004; 579:415–420.

9. Mekonnen D.W., Ludewig F. Phenotypic and chemotypic studies using Arabidopsis and yeast reveal that GHB converts to SSA and induce toxicity. Plant Mol. Biol. 2016; 91:429–440.

10. Simpson JP, Di Leo R, Dhanoa PK, Allan WL, Makhmoudova A, Clark SM, Hoover GJ, Mullen RT, Shelp BJ. Identification and characterization of a plastid-localized Arabidopsis glyoxylate reductase isoform: comparison with a cytosolic isoform and implications for cellular redox homeostasis and aldehyde detoxification. J Exp Bot. 2008; 59:2454–2554.

11. Hoover GJ, Van Cauwenberghe OR, Breitkreuz KE, Clark SM, Merrill AR, Shelp BJ. Characteristics of an Arabidopsis glyoxylate reductase: general biochemical properties and substrate specificity for the recombinant protein, and developmental expression and implications for glyoxylate and succinic semialdehyde metabolism in planta. Can J Bot. 2007; 85:883–895.

12. Ludewig F, Hueser A, Fromm H, Beauclair L, Bouche N (2008). Mutants of GABA transaminase (POP2) suppress the severe phenotype of succinic semialdehyde dehydrogenase (ssadh) mutants in Arabidopsis. PLoS One. 2008; 3(10):e3383.

13. van Cauwenberghe OR, Makhmoudova A, McLean MD, Clark S, Shelp BJ. Plant pyruvate-dependent γ-aminobutyrate transaminase: identification of an Arabidopsis cDNA and its expression inEscherichia coli. Can. J. of Bot. 2002; 80:933–941.

14. Clark SM, Di Leo R, Dhanoa PK, van Cauwengerghe OR, Mullen RT, Shelp BJ. Biochemical characterization, mitochondrial localization, expression, and potential functions for an Arabidopsis g-aminobutyrate transaminase that utilizes both pyruvate and glyoxylate. J Exp Bot. 2009; 60(6):1743–1757.

15. Poldrugo F, Addolorato G. The role of gamma-hydroxybutyric acid in the treatment of alcoholism: from animal to clinical studies. Alcohol & Alcoholism 1999; 34(1):15–24.

16. Jacobs C, Bojasch M, Mönch E, Rating D, Siemes H, Hanefeld F. Urinary excretion of gamma-hydroxybutyric acid in a patient with neurological abnormalities. The probability of the new inborn error of metabolism. Clin. Chim. Acta. 1981; 111:169–178.

17. Poirier Y, Antonenkov VD, Glumoff T, Hiltunen JK (2006). Peroxisomal beta-oxidation--a metabolic pathway with multiple functions. Biochim Biophys Acta. 2006; 1763(12):1413–26.

18. Masterson C., Wood C. Mitochondrial and peroxisomal beta-oxidation capacities of organs from a non-oilseed plant. Proc Biol Sci. 2001; 268(1479):1949–53.

19. Zolman BK, Silva ID, Bartel B. The Arabidopsis pxa1 mutant is defective in an ATP-binding cassette transporter-like protein required for peroxisomal fatty acid beta-oxidation. Plant Physiology. 2001; 127:1266–1278.

20. Footitt S, Cornah JE, Pracharoenwattana I, Bryce JH, Smith SM. The Arabidopsis 3 ketoacyl-CoA thiolase-2 (kat2-1) mutant exhibits increased flowering but reduced reproductive success. J. Exp. Bot. 2007; 58:2959–2968.

21. Jiang T, Zhang XF, Wang XF, Zhang DP. Arabidopsis 3-ketoacyl-CoA thiolase-2 (KAT2), an enzyme of fatty acid β-oxidation, is involved in ABA signal transduction. Plant Cell Physiol. 2011; 52(3):528–38.

